# The global impact of *Wolbachia* on mitochondrial diversity and evolution

**DOI:** 10.1101/160226

**Authors:** Marie Cariou, Laurent Duret, Sylvain Charlat

**Author notes:** Corresponding author. Sylvain Charlat.

## Abstract

The spread of maternally inherited microorganisms, such as *Wolbachia* bacteria, can induce indirect selective sweeps on host mitochondria, to which they are linked within the cytoplasm. The resulting reduction in effective population size might lead to smaller mitochondrial diversity and reduced efficiency of natural selection. Although suggested by a few case studies, the global consequences of this process on mitochondrial diversity and evolution remains to be assessed. Here we address this question using a mapping of Wolbachia acquisition / extinction events on a large mitochondrial DNA tree, including over 1,000 species. We show that the presence of Wolbachia is associated with a twofold reduction in silent mitochondrial polymorphism, and a 13% increase in non-synonymous substitution rates. These findings validate the conjecture that the widespread distribution of Wolbachia infections throughout arthropods impacts the effective population size of mitochondria. These effects might in part explain the disconnection between genetic diversity and demographic population size in mitochondria, and also fuel red-queen-like cytonuclear coevolution through the fixation of deleterious mitochondrial alleles.

## Introduction

Variations in population size have deep consequences on molecular evolution: small populations harbor fewer polymorphic sites and accumulate deleterious mutations at faster rates because of the predominance of drift. However, other, non-demographic processes, such as intense episodes of selection, also affect genetic diversity and substitution rates, which tends to uncouple the true population size from its abstract genetic counterpart, the effective population size (Ne). Mitochondrial DNA (mtDNA), although commonly used as a genetic marker for a number of good reasons (notably, technical ease-of-use and high mutation rates) is notoriously subject to such disconnection between the true and effective population size (Hurst and Jiggins 2005; Bazin et al. 2006; Galtier et al. 2009). Here we test the hypothesis that *Wolbachia* bacteria might be part of the explanation.

These intracellular (and thus maternally inherited) symbionts display an impressive variety of effects that make them invasive (O’Neill et al. 1997; Werren et al. 2008; Martinez et al. 2014). They can kill male embryos or turn them to females, reallocating part or all of the reproductive efforts toward the transmitting sex. They can also impede the reproduction of uninfected females, using infected males as sterilizing weapons. *Wolbachia* also commonly provides protection against natural enemies such as viruses, and thus indiscriminately benefits individuals of both sexes. In any case, if the net fitness gain to the infected maternal lineage is sufficient, *Wolbachia* can increase in frequency and drag along the mitochondrial lineage with which it happens to be associated, because the two are genetically linked through maternal transmission (Turelli et al. 1992).

If *Wolbachia* is perfectly transmitted from mothers to offspring, this process ends with the fixation of both the symbiont and the associated mitochondria, erasing the pre-existing molecular diversity. If transmission is imperfect, *Wolbachia* does not get fixed, but reaches an equilibrium prevalence, where selection for the infected lineage is balanced by imperfect transmission (O’Neill et al. 1997). Interestingly, even in that case, the ancestral mitochondrial polymorphism is erased in the long run, because all uninfected lineages ultimately originate from infected mothers, through imperfect transmission (Turelli et al. 1992). Thus, depending on the stage of the infection, the reduction in polymorphism within a species should either affect only its infected portion, or also extend to the uninfected individuals if the transmission / selection balance has been reached.

A number of case studies have demonstrated that the spread of *Wolbachia* can indeed affect the mtDNA polymorphism (Turelli et al. 1992; Solignac et al. 1994; Ballard et al. 1996; Jiggins 2003; Charlat et al. 2009; Graham and Wilson 2012; Obbard et al. 2012; Richardson et al. 2012; Schuler et al. 2016). Shoemaker (2004) also provided evidence for an elevated non-synonymous substitution rate in an infected *Drosophila* species compared to its uninfected sister species, making *Wolbachia* and reduction in Ne a very plausible explanation. Here we assess the generality of these effects using the SymbioCode system, a sample of more than one thousand Arthropod species collected in four Polynesian islands (Ramage et al. 2017). We map the previously inferred evolutionary history of *Wolbachia* acquisitions (Bailly-Bechet et al. 2017) on the mtDNA tree, and show that these symbionts affect both the mtDNA polymorphism and substitution rates. These results attest the marked global effect of *Wolbachia* infections on mitochondrial diversity and evolution.

## Material and Methods

### Dataset

The present analysis is based on the SymbioCode dataset which has been described in details elsewhere (Bailly-Bechet et al. 2017; Ramage et al. 2017) (dx.doi.org/10.5883/DS-SYMC). In brief 10,929 arthropod specimens were collected in four islands of the Society Archipelago, and sorted into morpho-species. DNA barcodes, that is, a 658 bp fragment of the CO1 mitochondrial gene, were obtained by Sanger sequencing from 3,627 specimens, spanning most of the taxonomic and geographic diversity of the initial sample (GenBank ids: KX051578 - KX055204). DNA barcodes clustered into 1,110 species-like groups or Operational Taxonomic Units (OTUs) covering 26 orders, the most species-rich being Diptera (306 OTUs), Lepidoptera (222), Hymenoptera (171), Hemiptera (132) and Coleoptera (106). The presence of *Wolbachia* was ascertained by a double PCR assay (Simões et al. 2011) in 32% of the specimens, and a standard *Wolbachia* marker, the fbpA gene, was directly sequenced from PCR products using Sanger sequencing (Bailly-Bechet et al. 2017). Specimens carrying uncharacterized *Wolbachia* strains (detected by PCR but not successfully sequenced) were excluded from the subsequent analysis, where sequence data was required. To eliminate possible cases of transient infections or artificial contaminations, we also filtered out OTUs carrying a single infected specimen from the present analysis.

A co-phylogenetic analysis based on the tree reconciliation program ALE (Szöllősi et al. 2013a,b) allowed us to map 1,000 plausible scenarios of *Wolbachia* acquisitions on the host mtDNA tree, sampled according to their likelihood (Bailly-Bechet et al. 2017). From these, we estimated the probability that any branch was infected as the proportion of scenarios where this branch was found infected. Notably, this proportion might differ between the start and the end of a branch (if infection was lost or acquired on this branch); we thus used the average of the two proportions as a measure of the infection probability.

### Nucleotide diversity and dN/dS estimations

Within each OTU, the silent nucleotide diversity (*π_s_*) was approximated as the mean of raw genetic distances between all specimens at the 3^rd^ position of codons in the CO1 gene using the R function *nuc.div* (PEGAS, Paradis 2010).

Synonymous (dS) and non-synonymous (dN) substitution rates were estimated on each branch of the host tree using a substitution mapping approach. In a first step, we estimated the ω parameter (ω=dN/dS) by maximum likelihood, using a homogeneous model, i.e. assuming a single ω parameter over the entire tree. In a second step, synonymous and non-synonymous substitutions are mapped on the tree to compute branch-specific values of ω. This approach was shown to be more accurate than a full maximum likelihood approach, where one would aim at estimating ω separately for each branch (O’Brien et al. 2009; Romiguier et al. 2012).

The maximum likelihood CO1 tree (Bailly-Bechet et al. 2017) was split in 18 clades of computationally manageable size. For each clade, this tree and the corresponding CO1 alignment were used to optimize an homogeneous substitution model by maximum likelihood (Yang and Nielsen 1998) with the program *bppml* (Bio++ *Maximum Likelihood*) (Dutheil and Boussau 2008). The model included the following parameters: equilibrium base composition, transition/transversion equilibrium ratio (κ=ts/tv), position specific base compositions at the root, and a single ω. *MapNH* (from the *TestNH* package) (Dutheil et al. 2012) was then used to count the number of synonymous and non-synonymous substitutions, as well as the number of synonymous and non-synonymous sites, on each branch of the trees. For each branch, the non-synonymous and synonymous substitution rates (dN and dS) are the ratio between the number of substitutions and the number of sites.

To compare ω values of infected and uninfected branches, we pooled estimates from all branches within small clades rather than use single branch estimates, for the two following reasons. First, dN and dS are inaccurately estimated on each specific branch, so that a pool of branches produces more robust estimates. Second, there is some uncertainty in the infection status of each branch, captured in our analysis by the estimated probability of infection. When this probability is not 0 or 1, a specific branch cannot be assigned to the uninfected or infected categories. However, by combining several branches, we can produce pooled estimates of the “infected” and “uninfected” ω by weighting data from every branch using its probability of infection. We used the mtDNA phylogeny to define 536 clades including specimens distant by no more than 0.2 substitutions per CO1 site (using branch length as a distance measure), and within each clade, we calculated the infected and uninfected ω as follows:

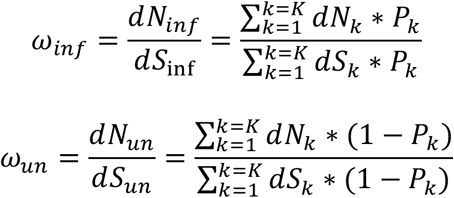

Where *P_k_* denotes the probability that branch *k* is infected and *d N_k_* and *d S_k_* denote the nonsynonymous and synonymous substitution rates on branch *k* (among K branches). This calculation ensures that the weight of each branch is proportional to its length and the level of confidence in its infection status. We limited this calculation to clades where ω_inf_ or ω_un_ could be estimated from sufficient data, that is, from branches summing to at least 2% in dS. The infected and uninfected ω values were thus estimated in 178 and 216 clades, respectively, with 176 clades providing estimates for both categories.

### Comparative analysis

Phylogenetic inertia can produce strong but spurious correlations between variables, or on the contrary blur correlations between causally linked variables (Felsenstein 1985). To test the effect of *Wolbachia* on either nucleotide diversity or substitution patterns, we controlled for this effect using paired comparisons. We used the above-defined 536 clades including specimens distant by no more than 0.2 substitutions per CO1 site. Within each clade including both infected and uninfected individuals, we then calculated the statistic of interest (π or ω) for the infected and uninfected categories. For the polymorphism analysis, these were simply the mean p values of each category of taxa. For the dN/dS analysis, the ω_inf_ and ω_un_ were computed as detailed above. For both analyses, we used Wilcoxon paired signed rank tests to assess differences between the infected and uninfected categories.

## Results

### Does *Wolbachia* reduce mitochondrial polymorphism?

To assess the effect of *Wolbachia*-induced sweeps, we used sequences of the CO1 gene to compare the silent mitochondrial polymorphism of 134 infected and 241 uninfected arthropod species, collected in French Polynesia as part of the SymbioCode project (Bailly-Bechet et al. 2017; Ramage et al. 2017). Despite a trend in the expected direction, this global comparison did not reveal a significant reduction of polymorphism linked with the presence of *Wolbachia* (figure 1a; mean uninfected *π_s_*=0.54%; mean infected *π_s_*=0.45%; Wilcoxon rank sum test, W=16947, p-value=0.38). However, this comparison can be confounded by background variation in mutation rates, census population size, or any other factor affecting polymorphism and varying across Arthropod clades. To control for these effects, we used a paired comparison on a subset of the data: 54 infected and 138 uninfected species distributed across 18 small clades (of maximum 20% CO1 divergence) including both categories. This more sensitive approach validates the hypothesis that *Wolbachia* reduces the mitochondrial polymorphism (Figure 1b, Wilcoxon paired test, V=28, p=0.04), with an overall twofold reduction associated with the presence of *Wolbachia* (mean uninfected *π_s_*=1.1%; mean infected *π_s_*=0.51%; n=18 clades). Importantly, this effect is visible in clades from several Arthropod orders (figure 1b), although the signal is necessarily less clear in those harboring a very low polymorphism.

**Fig. 1.**
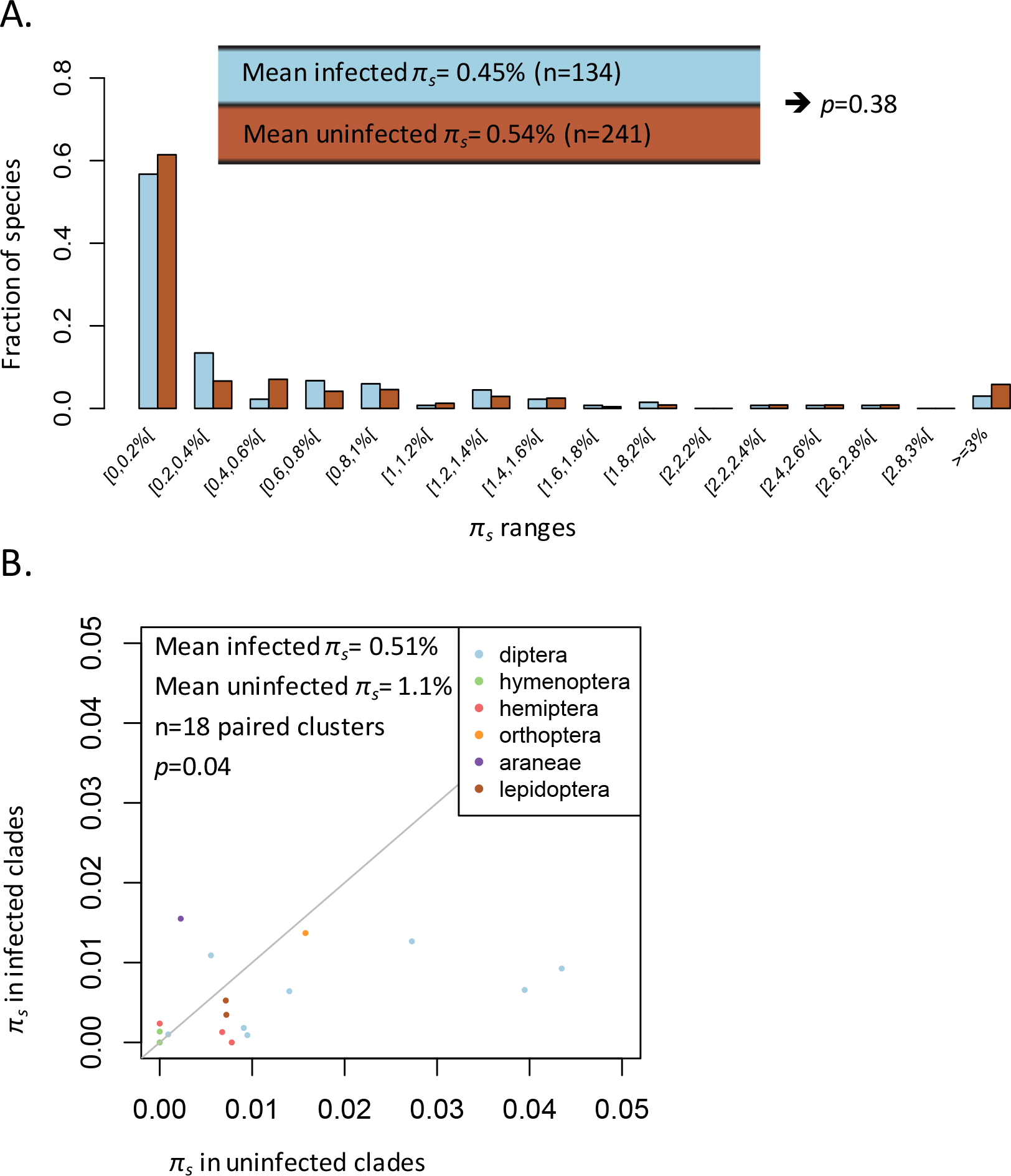
The effect of *Wolbachia* on silent nucleotidic diversity. A: distribution of *π_s_* in all infected and uninfected species (in blue and brown, respectively). B: paired comparison, plotting the infected versus uninfected mean *π_s_* computed across 18 clades carrying both categories of species.

### Does *Wolbachia* reduce purifying selection efficiency?

The spread of *Wolbachia* temporarily reduces the mitochondrial effective population size, which produces the above-documented reduction in polymorphism. But are these sweeps frequent and intense enough to also affect substitution patterns, that is, to increase the rate of fixation of non-synonymous mutations, most of which would otherwise be prevented by purifying selection? We tested this hypothesis by comparing ω, i.e. the ratio between nonsynonymous and synonymous substitution rates (dN/dS) among infected and uninfected lineages.

To this end, we used the output of a *Wolbachia* / mtDNA cophylogenetic analysis to estimate the probability that *Wolbachia* was present on each branch of the CO1 tree (Bailly-Bechet et al. 2017) and also estimated the number of synonymous and non-synonymous substitutions for each branch. To reduce the uncertainty in our analysis, we did not directly use branch specific estimates, but rather pooled the information from closely related branches, that is, branches belonging to the same clade of maximum 20% CO1 divergence. We further selected the pooled estimates that were based on sufficient total branch length (total dS > 2%). The final dataset thus includes 178 estimates for the “infected ω”, and 216 estimates for the “uninfected ω”, distributed across 218 clades of maximum 20% CO1 divergence.

Regardless of the presence of *Wolbachia*, we found that non-synonymous substitution rates are very low in all lineages, reaching about 1% of the synonymous substitution rate, in line with strong purifying selection acting on the mitochondrial genome CO1 gene (James et al. 2016). To assess the effect of *Wolbachia* on the efficiency of selection, we first compared the infected and uninfected ω values using a global, non-paired approach (figure 2a). Although the average infected ω is slightly larger than the average uninfected ω, this nonpaired test does not reveal a significant difference (mean ω_inf_=0.0086, n=178; mean ω_un_=0.0077, n=216; Wilcoxon rank sum test, W=17530, p-value=0.13). To better control for the effect of background variations in ω, we selected the 176 clades where both the infected and uninfected ω values could be computed and compared. This paired approach indicates a significant difference (Wilcoxon paired test, V= 5685, p=0.003), with a 13% increase in ω associated with the presence of *Wolbachia* (mean ω_inf_=0.0087, mean ω_un_=0.0077).

**Fig. 2.**
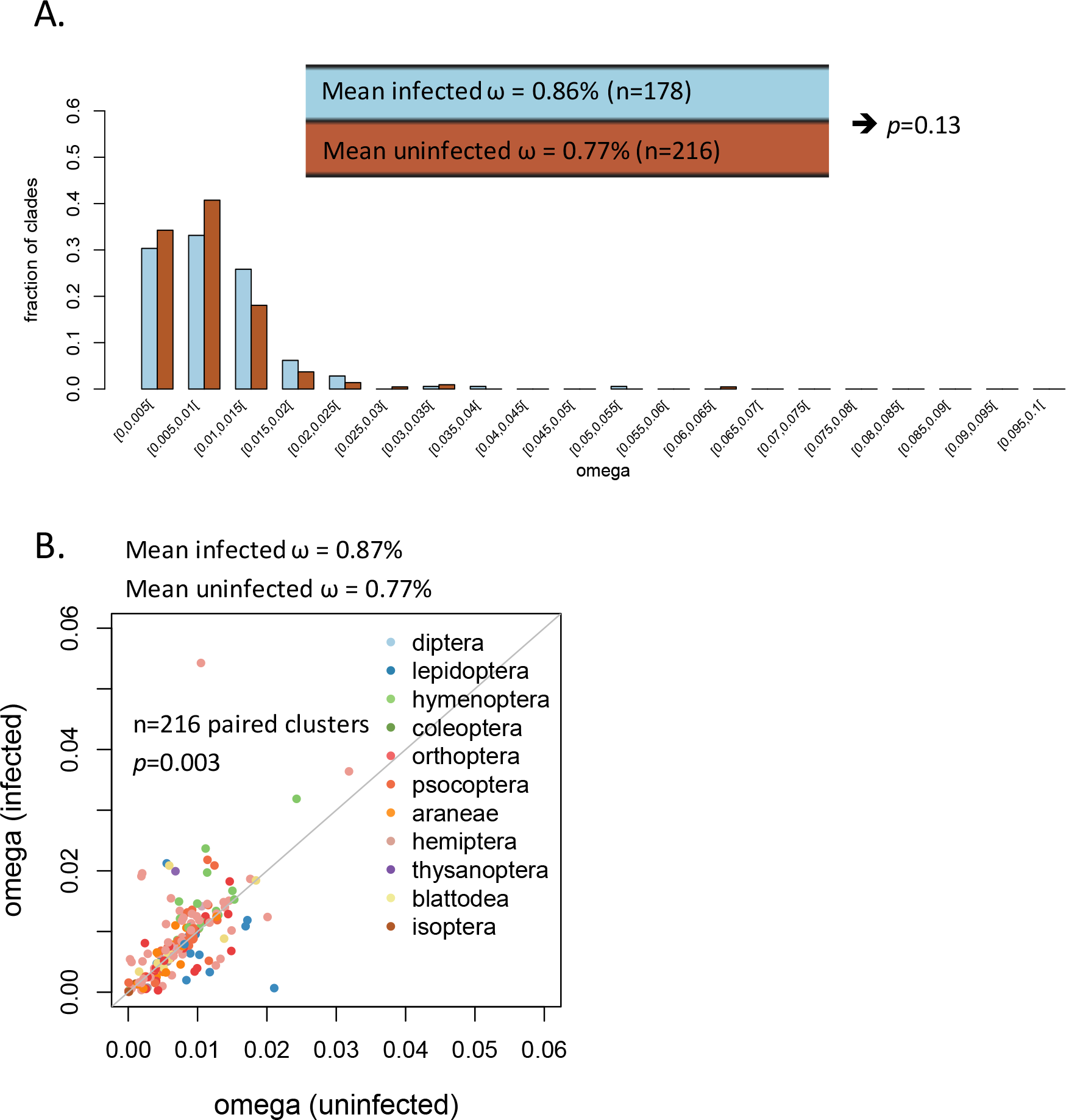
The effect of *Wolbachia* on non-synonymous substitution rates, standardized by synonymous substitution rates (ω=dN/dS). A: distribution of ω_inf_ or ω_un_ (in blue and brown, respectively) in all clades where at least one can be computed. B: paired comparison, plotting ω_inf_ versus ω_un_ across 176 clades where both ω_inf_ or ω_un_ were computed.

## Discussion

Theory suggests that *Wolbachia* infections should reduce the mitochondrial effective population size, while case studies indicate this can occasionally impact the polymorphism (Turelli et al. 1992; Solignac et al. 1994; Ballard et al. 1996; Johnstone and Hurst 1996b; Jiggins 2003; Charlat et al. 2009; Graham and Wilson 2012; Obbard et al. 2012; Richardson et al. 2012; Schuler et al. 2016) and possibly the efficacy of purifying selection (Shoemaker et al. 2004). The comparative approach used here suggests these effects are strong enough to globally affect the polymorphism and the molecular evolution of mitochondria. The presence of *Wolbachia* appears to be associated with a twofold reduction in polymorphism, and a 13% increase in non-synonymous substitution rate. We also note that other factors (e.g. variation in mutation rates) introduce variations in the silent mitochondrial polymorphism, so that the effects of *Wolbachia* are only detected with a paired approach, where we compare closely related infected and uninfected species or branches. This means that *Wolbachia* is one among several forces shaping the mitochondrial polymorphism and substitutions patterns across arthropods.

The observed reduction in polymorphism in *Wolbachia* infected species supports the hypothesis that natural selection acting on the symbiont has produced recent reductions in mitochondrial effective population size in many species. Under this view, several more specific and non-mutually exclusive hypotheses can be distinguished. First, and most obvious, it might be that the recent reduction in Ne was caused by the recent spread of new *Wolbachia* infections. A second possibility is that ancient infections are subject to recurrent selective sweeps, associated with repeated episodes of *Wolbachia* adaptive evolution within its host, which would reduce Ne beyond the initial invasion phase. Finally, a long-term reduction in Ne could also occur if the equilibrium prevalence is low. Indeed, the uninfected part of the population is an evolutionary dead end (that does not contribute to mitochondrial Ne), so that a low equilibrium prevalence can in principle maintain an abnormally low polymorphism in the long run (Johnstone and Hurst 1996a). Recent estimates of the *Wolbachia* turnover suggest that most infections have been acquired during the last few million years (Bailly-Bechet et al. 2017), a time frame that does not rule out any of the above explanations.

Interestingly we observed in our dataset that in species where both infected and uninfected lineages coexist, the mitochondrial polymorphism tends to be smaller in the infected portion than in the uninfected portion (*π_s_*=0.35% versus 0.53%). Theory and case studies indicate that once a maternally inherited symbiont has reached its equilibrium prevalence, the reduction in polymorphism also affects the uninfected part of the population, because this part is only maintained through the loss of infection from the infected lineage (Turelli et al. 1992; Solignac et al. 1994; Richardson et al. 2012). Our results indicate that infected species as a whole (including uninfected lineages) have a lower mitochondrial polymorphism than infected ones (0.51% versus 1.1%), suggesting the equilibrium prevalence has been reached in many cases, but also that the infected portion of infected species harbor an even lower polymorphism (0.35%), suggesting the equilibrium infection prevalence has not yet been reached in a substantial proportion of species. In this context, we also note that species carrying only one infected specimen were removed from the analysis; this allows us to eliminate natural or artificial *Wolbachia* DNA contaminations, but also tends to exclude infected species with low *Wolbachia* prevalence.

While a reduction of polymorphism provides hints on recent selective sweeps, the ω (i.e. dN/dS) ratio integrates all substitutions having occurred over long periods, and can thus reveal long-term variations in Ne. We found a 13% increase in ω associated with the presence of *Wolbachia* on branches of the CO1 tree. We can use previous estimation of the distribution of fitness effects (DFE) of non-synonymous mutations in mitochondria (James et al. 2016) to evaluate the magnitude of change in Ne that would be compatible with our observations. James et al. (2016) estimated the shape of the DFE from over 500 animal species. The shape parameter of this distribution provides a simple relationship between the proportion of nonsynonymous mutations becoming effectively neutral (Ohta 1977; Kimura 1979; Welch et al. 2008) when Ne is reduced: x^-λ^ =p (where x is the factor of change in Ne, λ is the shape parameter of the DFE, and p is the proportion of effectively neutral mutations). From this, we can derive an estimation of x, knowing p and λ: x=exp^-(log(p)/λ)^. We estimated that omega is 1.13 times larger in *Wolbachia* infected lineages. Assuming the majority of non-synonymous substitutions are deleterious, this means a 1.13 increase in the proportion of effectively neutral mutations. In other words, we estimate p=1.13. James et al (2016) estimate a global λ of 0.44, so that x=exp^-(log(1.13)/0.44)^=0.76. Thus, we estimate that Ne is reduced by 24% in lineages where *Wolbachia* is present, which, considering the various sources of uncertainty, is not incompatible with the 50% reduction in Ne estimated from the silent polymorphism data.

The global effect of *Wolbachia* on mitochondrial polymorphism and evolution argues against the view that *Wolbachia* might be frequently transmitted between different maternal lineages within species, either through occasional paternal transmission (Hoffmann et al. 1990), or horizontally transfer *sensu stricto* (Huigens et al. 2000, 2004). Thus, while non vertical transmission is known to occur and underlies the global distribution of this symbiont (Werren and Windsor 2000; Engelstädter and Hurst 2006; Zug et al. 2012; Bailly-Bechet et al. 2017), its rate appears too low to break the genetic linkage between *Wolbachia* and mitochondria within species.

Some central features of mitochondrial evolution should be revisited in the light of our findings. Notably, it has been shown that mitochondrial polymorphism is often disconnected from the true population size (Hurst and Jiggins 2005; Bazin et al. 2006; Galtier et al. 2009). Our results suggest that *Wolbachia* is likely part of the explanation. Evidence is also accumulating that coevolution between mitochondrial and nuclear genes often produces incompatibilities between recently isolated populations, thus contributing to the evolution of reproductive barriers (Burton et al. 2013; Chou and Leu 2015; Hill 2016). Specifically, it is hypothesized that mitochondrial properties (high mutation rate, maternal inheritance and lack of recombination) are responsible for the fixation of deleterious or selfish alleles, producing a red-queen-like cytonuclear coevolution. Under this view, mitochondria would represent an Achilles– heel for adaptive evolution, driving compensatory evolution in the nucleus. Our results suggest that the genetic linkage between mitochondria and widespread invasive cytoplasmic elements can exacerbate this process.

## Acknowledgments

We are very grateful to Laurent Gueguen for advice and assistance on the dN/dS analysis, to Nicolas Lartillot for helpful discussions on the comparative analysis, and to Greg Hurst and Fabrice Vavre for commenting an earlier version of this manuscript. This work was supported by the Centre National de la Recherche Scientifique (CNRS) (ATIP grant SymbioCode to S.C.) and benefitted from the computing facilities of the CC LBBE/PRABI.

## References

Bailly-Bechet, M., P. Simoes, G. Szöllősi, G. Mialdea, M.-F. Sagot, and S. Charlat. 2017. How long does Wolbachia remain on board? Mol. Biol. Evol. in press.

Ballard, J. W. O., J. Hatzidakis, T. L. Karr, and M. Kreitmant. 1996. Reduced variation in Drosophila simulans mitochondrial DNA. Genetics 144:1519–1528.

Bazin, E., S. Glemin, and N. Galtier. 2006. Population size does not influence mitochondrial genetic diversity in Animals. Science (80-.). 312:570–573.

Burton, R. S., R. J. Pereira, and F. S. Barreto. 2013. Cytonuclear Genomic Interactions and Hybrid Breakdown. Annu. Rev. Ecol. Evol. Syst. 44:281–302.

Charlat, S., A. Duplouy, E. A. Hornett, E. A. Dyson, N. Davies, G. K. Roderick, N. Wedell, and G. D. Hurst. 2009. The joint evolutionary histories of Wolbachia and mitochondria in Hypolimnas bolina. BMC Evol Biol 9:64.

Chou, J. Y., and J. Y. Leu. 2015. The Red Queen in mitochondria: Cyto-nuclear co-evolution, hybrid breakdown and human disease. Front. Genet. 6:1–8.

Dutheil, J., and B. Boussau. 2008. Non-homogeneous models of sequence evolution in the Bio++ suite of libraries and programs. BMC Evol. Biol. 8:255.

Dutheil, J. Y., N. Galtier, J. Romiguier, E. J. P. Douzery, V. Ranwez, and B. Boussau. 2012. Efficient selection of branch-specific models of sequence evolution. Mol. Biol. Evol. 29:1861–1874.

Engelstädter, J., and G. D. D. Hurst. 2006. The dynamics of parasite incidence across host species. Evol. Ecol. 20:603–616.

Felsenstein, J. 1985. Phylogenies and the comparative method. Am. Nat. 125:1–15.

Galtier, N., B. Nabholz, S. Glémin, and G. D. D. Hurst. 2009. Mitochondrial DNA as a marker of molecular diversity: a reappraisal. Mol. Ecol. 18:4541–4550

Graham, R. I., and K. Wilson. 2012. Male-killing Wolbachia and mitochondrial selective sweep in a migratory African insect. 1–11.

Hill, G. E. 2016. Mitonuclear coevolution as the genesis of speciation and the mitochondrial DNA barcode gap. Ecol. Evol. 6:5831–5842.

Hoffmann, A. A., M. Turelli, and L. G. Harshman. 1990. Factors affecting the distribution of cytoplasmic incompatibility in Drosophila simulans. Genetics 126:933–948.

Huigens, M. E., R. P. de Almeida, P. A. Boons, R. F. Luck, and R. Stouthamer. 2004. Natural interspecific and intraspecific horizontal transfer of parthenogenesis-inducing Wolbachia in Trichogramma wasps. Proc Biol Sci 271:509–515.

Huigens, M. E., R. F. Luck, R. H. Klaassen, M. F. Maas, M. J. Timmermans, and R. Stouthamer. 2000. Infectious parthenogenesis. Nature 405:178–179.

Hurst, G. D. D., and F. M. Jiggins. 2005. Problems with mitochondrial DNA as a marker in population, phylogeographic and phylogenetic studies : the effects of inherited symbionts. Proc. Natl. Acad. Sci. U. S. A. 272:1525–1534.

James, J. E., G. Piganeau, and A. Eyre-Walker. 2016. The rate of adaptive evolution in animal mitochondria. Mol. Ecol. 25:67–78.

Jiggins, F. M. 2003. Male-Killing Wolbachia and Mitochondrial DNA. Selective sweeps, hybrid introgression and parasite population dynamics. Genetics 164:5–12.

Johnstone, R. A., and G. D. D. Hurst. 1996a. Maternally inherited male-killing microorganisms may confound interpretation of mtDNA variation in insects. Biol. J. Linn. Soc. 53:453–470.

Johnstone, R. A., and G. D. D. Hurst. 1996b. Maternally inherited malelkilling microorganisms may confound interpretation of mitochondrial DNA variability. Biol. J. Linn. Soc. 58:453–470.

Kimura, M. 1979. Model of effectively neutral mutations in which selective constraint is incorporated. 76:3440–3444.

Martinez, J., B. Longdon, S. Bauer, Y. S. Chan, W. J. Miller, K. Bourtzis, L. Teixeira, and F. M. Jiggins. 2014. Symbionts Commonly Provide Broad Spectrum Resistance to Viruses in Insects: A Comparative Analysis of Wolbachia Strains. PLoS Pathog. 10:e1004369.

O’Brien, J. D., V. N. Minin, and M. A. Suchard. 2009. Learning to count: Robust estimates for labeled distances between molecular sequences. Mol. Biol. Evol. 26:801–814.

O’Neill, S. L., A. A. Hoffmann, and J. H. Werren. 1997. Influential Passengers : Inherited Microorganisms and Arthropod Reproduction. Oxford University Press, Oxford.

Obbard, D. J., J. MacLennan, K. W. Kim, A. Rambaut, P. M. O’Grady, and F. M. Jiggins. 2012. Estimating divergence dates and substitution rates in the drosophila phylogeny. Mol. Biol. Evol. 29:3459–3473.

Ohta, T. 1977. Extension to the neutral mutation random drift hypothesis. Pp. 148–167 in M. Kimura, ed. Molecular evolution and polymorphism: Proceedings of the Second Taniguchi International Symposium on Biophysics.

Paradis, E. 2010. pegas : an R package for population genetics with an integrated - modular approach. Bioinformatics 26:419–420.

Ramage, T., P. Martins-Simoes, G. Mialdea, R. Allemand, A. M. R. Duplouy, P. Rousse, N. Davies, G. K. Roderick, and S. Charlat. 2017. A DNA barcode-based survey of terrestrial arthropods in the Society Islands of French Polynesia: host diversity within the SymbioCode project. Eur. J. Taxon. 272:1–13.

Richardson, M. F., L. A. Weinert, J. J. Welch, R. S. Linheiro, M. M. Magwire, F. M. Jiggins, and C. M. Bergman. 2012. Population Genomics of the Wolbachia Endosymbiont in Drosophila melanogaster. PLoS Genet. 8:e1003129. Public Library of Science.

Romiguier, J., E. Figuet, N. Galtier, E. J. P. Douzery, B. Boussau, J. Y. Dutheil, and V. Ranwez. 2012. Fast and robust characterization of time-heterogeneous sequence evolutionary processes using substitution mapping. PLoS One 7:e33852.

Schuler, H., K. Köpler, S. Daxböck-horvath, B. Rasool, S. Krumböck, D. Schwarz, T. S. Hoffmeister, B. C. Schlick-steiner, F. M. Steiner, A. Telschow, C. Stauffer, W. Arthofer, and M. Riegler. 2016. The hitchhiker’s guide to Europe : the infection dynamics of an ongoing Wolbachia invasion and mitochondrial selective sweep in Rhagoletis cerasi. 1595–1609.

Shoemaker, D. D., K. A. Dyer, M. Ahrens, K. McAbee, and J. Jaenike. 2004. Decreased diversity but increased substitution rate in host mtDNA as a consequence of Wolbachia endosymbiont infection. Genetics 168:2049–2058.

Simões, P. M., G. Mialdea, D. Reiss, M.-F. Sagot, and S. Charlat. 2011. Wolbachia detection: an assessment of standard PCR protocols. Mol. Ecol. Resour. 11:567–72.

Solignac, M., D. Vautrin, and F. Rousset. 1994. Widespread occurence of the proteobacteria Wolbachia and partial incompatibility in Drosophila melanogaster. Comptes Rendus l’Academie des Sci. Paris, Série III 317:461–470.

Szöllősi, G. J., W. Rosikiewicz, B. Boussau, E. Tannier, and V. Daubin. 2013a. Efficient exploration of the space of reconciled gene trees. Syst. Biol. 62:901–912.

Szöllősi, G. J., E. Tannier, N. Lartillot, and V. Daubin. 2013b. Lateral gene transfer from the dead. Syst. Biol. 62:386–397.

Turelli, M., A. Hoffmann, and S. McKechnie. 1992. Dynamics of cytoplasmic incompatibility and mtDNA variation in Natural Drosophila simulans populations. Genetics 132:713–723.

Welch, J. J., A. Eyre-walker, and D. Waxman. 2008. Divergence and Polymorphism Under the Nearly Neutral Theory of Molecular Evolution. J. Mol. Evol. 67:418–426.

Werren, J. H., L. Baldo, and M. E. Clark. 2008. Wolbachia: master manipulators of invertebrate biology. Nat. Rev. Microbiol. 6:741–751.

Werren, J. H., and D. M. Windsor. 2000. Wolbachia infection frequencies in insects: evidence of a global equilibrium? Proc Biol Sci 267:1277–1285.

Yang, Z., and R. Nielsen. 1998. Synonymous and Nonsynonymous Rate Variation in Nuclear Genes of Mammals. J. Mol. Evol. 46:409–418.

Zug, R., A. Koehncke, and P. Hammerstein. 2012. Epidemiology in evolutionary time : the case of Wolbachia horizontal transmission between arthropod host species. 1–12.

